# Alpha-synuclein fibrils induce autophagy in microglial cells as a consequence of lysosomal damage

**DOI:** 10.1101/169375

**Authors:** Claudio Bussi, Javier M. Peralta Ramos, Daniela S. Arroyo, Jose I. Gallea, Paolo Ronchi, Androniki Kolovou, Ji M. Wang, Oliver Florey, Maria S. Celej, Yannick Schwab, Nicholas T. Ktistakis, Pablo Iribarren

## Abstract

Autophagy is a constitutive lysosomal catabolic pathway that degrades damaged organelles and protein aggregates. In Parkinson’s disease, the synaptic protein alpha-synuclein (AS) accumulates in neuronal cell bodies and axons. Recent studies indicate that aggregation-prone proteins can spread to other brain cells - such as glia - contributing to progressive deterioration.

Although autophagic dysfunction and protein aggregation have been linked to several neurodegenerative disorders, exact mechanisms are not clear and most work was done in neurons and not on microglial cells.

Here we report that AS fibrils but not monomers induce lysosomal damage and autophagy in microglial cells and we extensively characterized the dynamics of this response by both live-cell imaging and correlative light-electron microscopy (CLEM). In addition, we found that autophagy inhibition in these cells impairs mitochondrial quality and leads to microglial cell death. We propose that AS accumulation in lysosomes leads to lysosomal damage, which in turn activates canonical autophagy as a rescue mechanism.

Our results provide novel findings about the interaction between AS and the autophagy pathway in microglial cells, which may be important for targeting protein misfolding-associated neurodegenerative diseases.

## INTRODUCTION

Neurodegenerative diseases are characterised by common cellular and molecular mechanisms including protein aggregation and inclusion body formation that result in toxicity and neuronal cell death [42].

Autophagy dysfunction in the central nervous system (CNS) has been shown to induce neurodegeneration even in the absence of any disease-associated mutant proteins. Mice deficient for *Atg5* (autophagy-related 5) develop progressive deficits in motor function that are accompanied by the accumulation of cytoplasmic inclusion bodies in neurons [26]. On the other hand, mice lacking *Atg7* specifically in the CNS showed behavioural defects, a reduction in coordinated movement and massive neuronal loss in the cerebral and cerebellar cortices [33].

Although latest developments reveal a crucial role for the autophagy pathway in neurodegenerative diseases [23], the precise mechanisms underlying these processes are poorly understood. Furthermore, most of the existing literature related to autophagy in the CNS focuses on neurons and the effects of the autophagy pathway and its modulation on microglial cells remain poorly understood. Microglia are resident macrophage cells in the CNS and have multiple functions, such as phagocytosis, production of growth factors and cytokines, and antigen presentation. The major function of microglia is to maintain homeostasis and normal function of the CNS, both during development and in response to CNS injury [56].

Canonical autophagy starts with the assembly of a pre-initiation complex consisting of ULK1, FIP200, and ATG13, which in turn leads to activation of the VPS34/Beclin-1 PI3K complex and then formation and extension of a double-membraned autophagosome around cellular contents by the lipidation of the autophagic protein light chain 3 (LC3), through the action of two ubiquitin-like conjugation systems. ULK1 is subject to regulatory phosphorylation by mTOR and AMPK, and this provides a means for the control of autophagy in response to nutrient status [36].

Lipidated LC3 was once thought to unambiguously distinguish autophagosomes from other cellular membranes. However, these past recent years, a non-canonical autophagy mechanism was reported in the literature depending on direct LC3 association with single limiting-membrane vacuoles and able to deliver the luminal content towards lysosomal degradation [45]. This unconventional pathway is known as LC3-associated phagocytosis (LAP) and it is involved in the maturation of single-membrane phagosomes and subsequent killing of ingested pathogens by phagocytes. LAP is initiated following recognition of pathogens by pattern recognition receptors and leads to the recruitment of LC3 into the phagosomal membrane [46].

Parkinson’s disease (PD) is a late-onset neurodegenerative disorder caused by degeneration of dopaminergic neurons in the substantia nigra. This pathology is characterized by the presence of intracellular inclusions named Lewy bodies, which contain the proteins AS and ubiquitin. AS is a presynaptic neuronal protein that is linked genetically and neuropathologically to PD. AS may contribute to PD pathogenesis by distinct mechanisms, but novel evidences suggest that its aberrant fibril conformations are the toxic species that mediate disruption of cellular homeostasis and neuronal death, through effects on various intracellular targets, including synaptic function [55]. Furthermore, secreted AS may exert deleterious effects on neighbouring neuronal and glial cells, including seeding of aggregation, thus contributing to disease propagation. Recent research suggests a complex role for microglia not only in PD but in other disorders involving AS aggregation, such as multiple system atrophy [58]. In addition, the novel concepts of AS being released in exosomes and uptaken by neighbouring cells, and their importance in disease progression, positions microglia as the main cell that can efficiently clear and handle AS [11, 41].

The pathogenic role of autophagy in PD was demonstrated by the finding that AS is degraded by macroautophagy and chaperone-mediated autophagy [64]. Emerging evidence has suggested that aberrant autophagy is one of the underlying mechanisms for hereditary forms of PD [31] and most of these studies showed that these genetic defects lead to autophagy impairment [52]. In addition, it has been shown that in degenerating neurons in PD brains, there is a breakdown of lysosomal membranes and mislocalization of lysosomal receptors and autophagy components [3, 29]. On the other hand, recent reports indicate that AS induce mitochondrial and lysosomal dysfunction and alters vesicular trafficking in PD, which may lead to AS accumulation [15, 48]. In this scenario the autophagy pathway plays a pivotal role as the main mechanism responsible for abnormal protein and organelle degradation [53].

Although significant progress has been done in unravelling the role and regulation of the autophagy machinery, its dysfunction in pathology as well as its dynamic changes in the disease progression remains largely unclear [42]. Further characterization of autophagy dynamics not only in neuronal but also in glial cells in combination with analysis of the ultrastructural details of the many novel organelles and mechanisms involved in specific subtypes of autophagy, for example, by discriminating autophagosomes from LAP, i.e. double-from single-limiting membrane of LC3-positive vesicles, may contribute for the development of novel therapeutic strategies in PD and other neurodegenerative disorders. In this report we investigated for the first time the effect of monomeric and fibrilar AS on the autophagy activity of microglial cells by live-cell imaging and EM techniques. We found that only fibrilar AS induces autophagy in microglial cells. Furthermore, we extensively characterized the dynamics of this response and we observed that activation of the autophagy pathway is concomitant to lysosomal damage. In addition, we analysed by high-precision and live CLEM experiments the ultrastructural morphology of the autophagic vesicles formed in AS stimulated cells. Moreover, we observed that autophagy inhibition led to mitochondrial quality impairment and microglial cell death after AS stimulation.

## MATERIALS AND METHODS

### Reagents

Dulbecco’s Modified Eagle Medium (DMEM), Opti-MEM, fetal bovine serum (FBS), penicillin, glutamine, G418, streptomycin, Silencer select FIP200 siRNA (4390771), Silencer select negative control 1 (4390843), Lipofectamine RNAiMAX (13778100) and LysoTracker Blue (L7525) were obtained from Thermo Fischer Scientific (Grand Island,
NY, USA). Antibodies used for western blotting were rabbit polyclonal anti-LC3 (Sigma, L7543) and mouse monoclonal anti β-actin (Cell Signaling, 8H10D10). Antibodies used for immunofluorescence were rabbit monoclonal anti-LC3 A/B (Cell Signaling, D3U4C), rat monoclonal anti LAMP-1 (Biolegend, 121601), mouse monoclonal anti-Galectin-3 (Biolegend, 125401), rabbit polyclonal anti-GALNS (GeneTex, 110237). PP242 (13643) and BafA1 (11038) were purchased from Cayman (Ann Arbor, MI, USA). Spautin-1 (SML0440) was purchased from Sigma. MitoSpy™ Green FM and MitoSpy™ Orange CMTMRos were obtained from Biolegend (San Diego, CA, USA). Magic Red™ Cathepsin-B Kit was purchased from Bio-Rad (Hercules, CA, USA). FITC Annexin V Apoptosis Detection Kit was purchased from Becton Dickinson (San Jose, CA, USA).

### Cell culture and transfections

The murine microglial cell line BV2 was a kind gift from Dr. Dennis J. Selkoe (Harvard Medical School, Center for Neurological Diseases, Brigham and Women’s Hospital, Boston, MA, USA). The cells were grown in DMEM supplemented with 10% heat-inactivated FCS, 2mM glutamine and 100 ug/ml streptomycin and maintained at 37°C and 5% CO2.

For stable GFP-LC3 expression, BV-2 cells were transduced with pBabe-GFP-LC3 retrovirus generated as previously described [22]. Cells were selected with 10μg/ml blasticidin for 4 days. BV2 stably transfected with ATG13 were cultured in media identical in composition to wild-type media, except for the addition of 400 μg/ml G418. BV2 cells were transfected with X-tremeGENE™ HP DNA Transfection Reagent (Roche, 06366236001) following manufacturer’s indications. CFP-LC3 plasmid was a kind gift from Dr. Tamotsu Yoshimori. Silencer Select FIP200 (Thermo Fischer, 4390771) and negative control siRNAs (Thermo Fischer, 4390843) solutions were prepared according to the manufacturer’s instructions and experiments were performed 48h after transfection. Lipofectamine RNAiMAX (Thermo Fischer, 13778100) was used as transfection reagent.

### Isolation of primary microglial cells from adult mice

After perfusion with PBS, brains from 6- to 8-wk-old mice C57BL/6J, (15 mice/group) were collected in DMEM, dispersed with scissors, resuspended in PBS containing 0.3% collagenase D (Roche, Indianapolis, IN, USA) and 10 mM HEPES buffer (Invitrogen, Carlsbad, CA, USA), and incubated 30 min at 37°C. Brain homogenates were then filtered in 70-_m-pore cell strainers (Becton Dickinson), centrifuged (7 min, 1500 rpm), washed, and resuspended in 70% isotonic Percoll (GE Healthcare, Fairfield, CT, USA). Cell suspension (3.5 ml) was transferred to 15-ml polypropylene conical tubes with 5 ml of 25% isotonic Percoll, which were sequentially layered on top with 3 ml of PBS. After centrifugation (30 min, 800 g, 4°C), the 70%:25% Percoll interphase layers were collected, and the cells were washed. Finally, the adherent cells, which contained 90% of CD11b+ cells, were cultured in DMEM supplemented with 10% heat-inactivated FCS, 2 mM glutamine, 100 U/ml penicillin, 100 ug/ml streptomycin, 100 ug/ml sodium pyruvate, and 10 mM HEPES buffer (Invitrogen). Microglial cells were washed with PBS and resuspended in medium containing 10% heat-inactivated FCS, alpha-synuclein, or other stimuli and then cultured for the indicated times at 37°C. Morphological changes were observed in a contrast phase microscope. Animal care was provided in accordance with the procedures outlined in the U.S. National Institutes of Health (NIH) Guide for the Care and Use of Laboratory Animals (Publication 86-23, 1985). The experimental protocols were approved by the Institutional Animal Care and Use Committee of Centro de Investigaciones en Bioquímica Clínica e Inmunología (CIBICI), Consejo Nacional de Investigaciones Científicas y Técnicas (CONICET). Our animal facility obtained NIH animal welfare assurance (assurance no. A5802-01, Office of Laboratory Animal Welfare, NIH, Bethesda, MD, USA).

### Cell culture treatments

BV2 cells were grown in six well plates to 65 to 70% confluence (for immunofluorescence) or 70 to 80% confluence (for western blotting) before treatments. Primary microglial cells were grown in chamber slides or 48 well plates to 65 to 70% confluence before treatments. AS fibrils and monomers were used at 5μM at indicated time points. PP242 was used at 1μM for 3h. Mammalian PI3KC3 was blocked by the addition of spautin-1 (10 μM) for 24h before stimulation. Autophagosome maturation was blocked using BAF, at final concentration of 200 nM, in normal growth medium for 1 h, unless otherwise stated.

### Preparation of monomeric and aggregated AS

Monomeric AS stock solutions were prepared in PBS buffer and 0.02% sodium azide and centrifuged (14100 g, 30 min) before use in order to remove possible aggregates. After that, solutions were sterilized by filtration (22 um pore size). Protein concentration was determined by absorbance using an ε275 of 5600 M−1 · cm−1. Fibrillation was achieved by incubating 400 μM monomeric AS stock solutions at 70°C and 800 rpm in a Thermomixer5436 (Eppendorf), conditions that lead to faster aggregation kinetics [10] Fibril formation was monitored using the ThioT (thioflavin T) fluorescence assay. Fibrils were isolated by three consecutive cycles of centrifugation (14000 g, 30 min) and resuspended in PBS buffer. Protein concentrations in monomeric units were determined by the absorbance of aliquots incubated in 6 M guanidinium chloride at 25°C for 24 h. Endotoxin levels were evaluated by Limulus Amebocyte Lysate (LAL) assay. Endotoxin content was lower than detection limit (<0,24 EU/mL). AS protein labelling: AS fibrils were conjugated by using Alexa Fluor 647 (Molecular probes, A2006) or Alexa Fluor 546 (Molecular Probes, A2004) NHS Ester dye according to manufacturer’s instructions.

#### Western immunoblotting

After treatment with AS fibrils or monomers, 2x106 BV2 microglial cells were harvested by centrifugation at indicated time points. After washing with PBS, cells were lysed using sample buffer. Subsequently, cells lysates were sonicated and boiled. Proteins were electrophoresed on 12% SDS-PAGE gel under reducing conditions and transferred on to Immun-Blot PVDF Membrane (Bio-Rad, Hercules, CA, USA). The membranes were blocked with 5% nonfat milk and 0.1% Tween-20 in TBS for 2h at room temperature and then were incubated with primary antibodies overnight at 4°C. Then, membranes were incubated with secondary antibodies (IRDye, LI-COR Biosciences, Lincoln, Nebraska, USA) for 1h and 30 min at RT and protein bands were detected with an Odyssey Infrared Imaging System (LI-COR Biosciences).

#### Immunofluorescence assays

Following the appropriated treatments BV2 cells grown on glass coverslips in 6-well plates, or primary microglial cells grown on Lab-Tek chamber slides (Thermo Fischer), were fixed with 100% ice-cold metanol for 10 min on ice. The cells were then blocked for a minimum of 1 h in 5% (w/v) BSA)/PBS before staining with the appropriate primary antibodies. After 3 rinses with PBS, samples were incubated with Alexa Fluor 488 or Alexa Fluor 546 secondary antibodies (Invitrogen) for 60 min. The slides were analyzed under a laser scanning confocal fluorescence microscope (Olympus FV1000; Olympus, Tokyo, Japan) or under a wide-field fluorescence microscope (Leica DMi8, Leica, Weitzlar, Germany). Galectin puncta assay: Staining for immunofluorescence and image analysis were done following a previously characterized protocol (Aits et al., 2015). Briefly BV2 or primary microglial cells were incubated in the presence or the absence of fibrilar AS and immunolabeled with a monoclonal rat anti-mouse galectin 3 antibody (Biolegend, 125402). Alternatively, immunostaining with a monoclonal rat anti-mouse galectin-3 Alexa Fluor 488 conjugated antibody (Biolegend, 125410) was also performed. After that, fluorescence images were acquired under a wide-field or laser scanning confocal fluorescence microscope and galectin-3 puncta formation was followed over time. Incubation for 2h at 37°C with a 500 μM solution of LLOME crystals was used as positive control of lysosomal damage. Spatial deconvolution, 3D surface-rendered images and 3D surface-rendered movies were carried out with SVI Huygens Software. Image processing and analysis were performed using FIJI software (NIH) (Schindelin, Arganda-Carreras et al., 2012).

#### Live-cell imaging

Live-cell imaging was performed in cells that had been plated on to 22-mm diameter coverslips (BDH) and transiently transfected with the relevant constructs. Individual coverslips were placed in an imaging chamber with 2 ml of medium and the appropriate treatment. LysoTracker Blue (1:10000; Thermo Fischer, L7525), where stated, was added to samples 30 min before imaging began. Samples were then placed in a Solent environment chamber (Solent Scientific, custom made) before mounting on the microscope, all at 37°C. Widefield imaging experiments were performed on a Nikon Ti-E-based system. The Nikon Ti-E-based system comprised a Nikon Ti-E microscope, 100x 1.4 N.A. objective (Nikon), SpecraX LED illuminator (Lumencor), 410/504/582/669-Di01 and Di01-R442/514/561 dichroic mirrors (Semrock), Hamamatsu Flash 4.0 sCMOS camera, emission filter wheel (Sutter Instruments) and was controlled using Nikon Elements software. Excitation and emission filters (all from Semrock) were as follows: CFP 434/17 (ex) 480/17 (em), GFP 480/10 (ex) 525/30 (em), mRFP 560/25 (ex) 607/36 (em).

#### Correlative Light-Electron Microscopy (CLEM) experiments

Widefield-electron microscopy correlation assays: BV2 GFP-LC3 cells were cultured on 3mm carbon-coated sapphire discs and stimulated with labelled AS (3uM) for 12h. High pressure freezing, ultrathin sectioning for electron microscopy, image acquisition and correlation analysis were done as described previously [37]. Briefly, after cellular stimulation, microglial cells were high pressure frozen using an Abra Fluid HPM-010 (Abra Fluid, Switzerland) and transferred to the automated freeze substitution apparatus (Leica EM AFS2) under liquid nitrogen. Semithick 300-nm sections were prepared using a Leica UC6/UF6 ultramicrotome and picked up on finder 200 mesh copper grids coated with carbon. After that, immunofluorescence images were acquired using an Olympus Scan^R microscope. Previous to the EM acquisition, colloidal gold particles, 10 nm in diameter, were placed on top of the sections to serve as fiducial markers for alignment of the tomograms. Tilt series were acquired on a Tecnai F30 microscope (FEI, Netherlands) operating at 300 kV with a OneView camera (Gatan Inc., USA) at a binned (2) pixel size of 1.25 nm using SerialEM [47]. Images were recorded at 1 degree intervals over a tilt range of +60 to -60 degrees. Electron and fluorescent images overlay were obtained by using ec-CLEM plugin [54] from ICY software [13] IMOD software package [34] was used to create 3D reconstructions from the tilt series and to create 3D models of the autophagosomes and membranes. LIVE-CLEM assays: BV2 GFP-LC3 cells were grown on a gridded MatTek (Ashland, MA, USA) and live-imaged by widefield microscopy using a Zeiss Celldiscoverer7 microscope, (Zeiss, Germany) after 12h of AS stimulation. A bright field image of the area containing the cell of interest was acquired at low magnification to visualize the grid and therefore precisely localize the position of the cell. Cells were fixed in 2.5% glutaraldehyde (GA, Electron Microscopy Sciences) in 0.1M cacodylate buffer immediately after detecting the event of interest. The subsequent EM processing steps (OSO4, UA, dehydration) were performed using a PELCO Biowave Pro microwave processor (Ted Pella, Inc.). After dehydration the coverslip was detached from the MatTek dish and put on an Epon-filled capsule. After polymerization, the area containing the cell of interest was retrieved by means of the grid coordinate system that remained impressed on the block surface. The blocks were then sectioned with a Leica UC7 ultramicrotome and 300nm sections were collected on formvar coated slot grids. Tilt series of the cell of interest were acquired with a FEI Tecnai F30 electron microscope. Tomogram reconstruction, segmentation and 3D rendering was carried out with the IMOD software package [34].

### Evaluation of mitochondrial quality and Cathepsin-B activity

BV2 microglial cells were left untreated or stimulated with fibrilar AS from 8 to 48h. After that, the cells were harvested, washed twice with fresh medium and incubated for 30min at 37°C in DMEM supplemented with 10% heat-inactivated FCS containing the following dyes: 100 nM MitoSpy Green FM (Biolegend, 424805) to measure mitochondrial mass, 100 nM MitoSpy Orange CMTMRos (Biolegend, 424803) to measure mitochondrial membrane potential and 26X Magic Red Cathepsin-B substrate (BioRad, ICT937) to measure Cathepsin-B activity. Cells were then washed and resuspended in 300uL of FACS buffer. Flow cytometric analysis were performed on a FACSCanto II cytometer (Becton Dickinson) using FCS De Novo Software.

### Evaluation of cell death by flow cytometry

BV2 microglial cells and primary microglial cells were washed twice with PBS and incubated with propidium iodide (PI) for 2 minutes in 300uL of FACS buffer. For annexin V (AnV) and PI dual staining, the cells were harvested, washed twice with binding buffer, and incubated with FITC-conjugated AnV and PI following manufacturer instructions (FITC Annexin V Apoptosis Detection Kit, Becton Dickinson). For Bcl-2, Bcl-xL and cleaved caspase-3 staining, BV2 cells were fixed and permeabilized by using Cytofix/Cytoperm kit (Becton Dickinson) and incubated with the monoclonal primary antibodies (Cell Signaling). After 1h, cells were washed and incubated for 30 min with an anti-rabbit Alexa Fluor 488 antibody. After that cells were washed with PBS and resuspended in 300uL of FACS buffer. Microglial cells were then analyzed by flow cytometry on a FACSCanto II cytometer (Becton Dickinson) using FCS De Novo Software.

### Statistical analyses

The results were analysed using one-way analysis of variance (ANOVA) model, as indicated for every experiment. GraphPad Prism 6.0 was used to carry out the computations for all analyses. Results represent mean±SEM of at least three experiments. Statistical significance was defined as p≤0.05.

## RESULTS

### Fibrilar but not monomeric AS induces autophagy in microglial cells

We first analysed the effect of different AS aggregation states on autophagy induction in microglial cells. We stimulated BV2 and primary microglial cells with both exogenous fibrilar and monomeric AS at different time points. Based on a recent report from our group, we selected 5 uM as protein concentration since no significant toxicity is observed either in BV2 or primary microglial cells after long-term cell culture [9]. Interestingly, we observed by immunofluorescence visualization (Fig. 1A) that only fibrilar, but not monomeric AS, increased the punctate localization of LC3. In addition, this response increased substantially after 12h of stimulation (Fig.1A). We did not find changes in the autophagic response after monomeric stimulation at all time-points studied.

**Figure 1.**
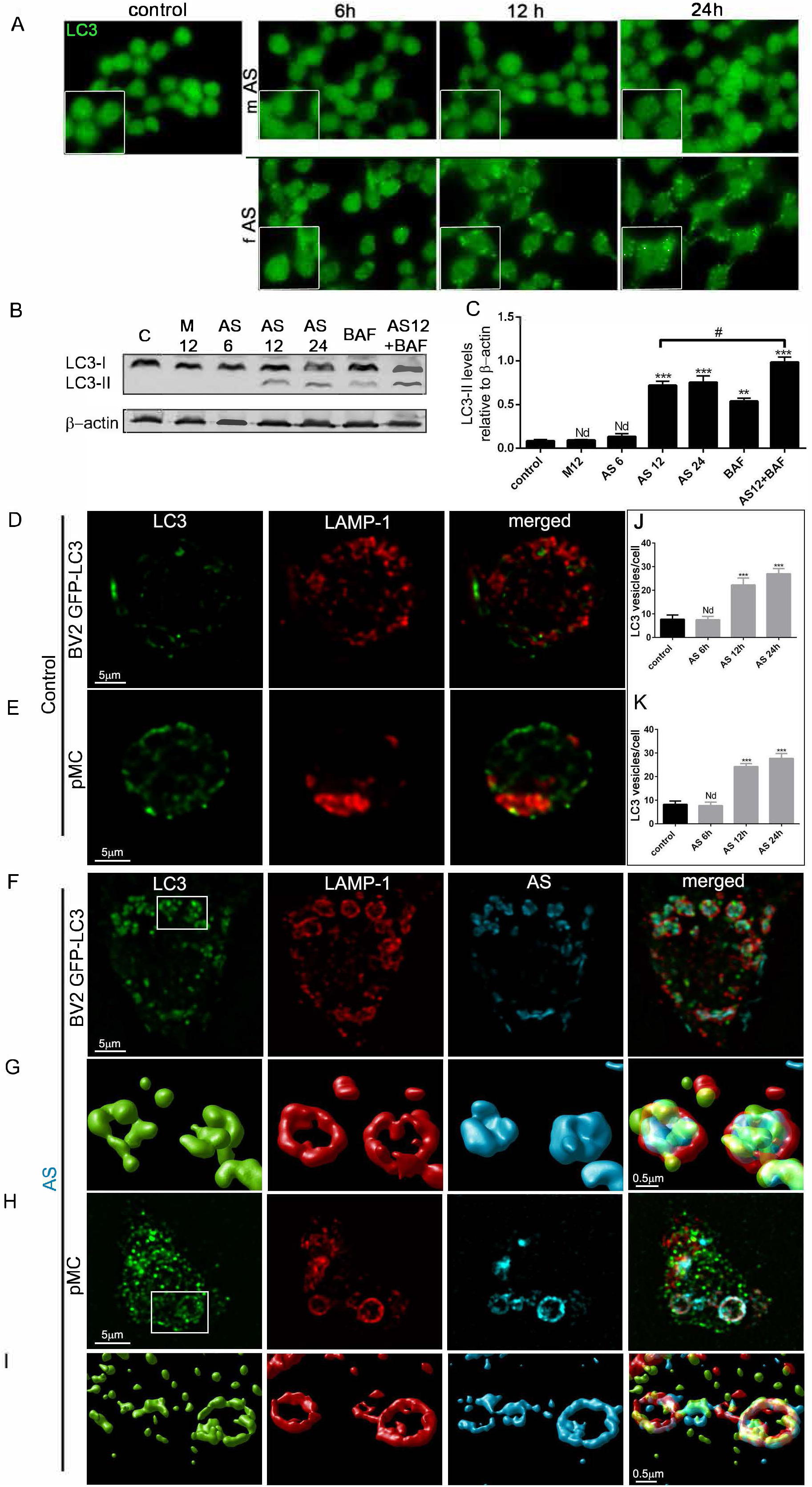
Alpha-synuclein induces autophagy in microglial cells. (A) BV2 microglial cells were left untreated or stimulated at different time points with AS monomers (m) or fibrils (f) at 5uM. Cells were then fixed and stained for LC3. (B) Cell lysates from BV2 cells cultured with AS fibrils or monomers (5 uM) were collected at different time points and LC3B and β-Actin protein levels were examined by Western immunoblotting. Bafilomycin A1 (BAF) was added for the last 3h. (C) Quantification of LC3-II from (D) relative to β-Actin by densitometry. BV2 GFP-LC3 cells (D, E) or primary microglial cells (F, H) were left untreated or stimulated with Alexa Fluor 647-labeled AS fibrils (5uM). After 12h cells were immunostained with anti-Lampl (red) antibody and primary microglial cells were also stained for LC3. Images shown are z-stack projections. (G) and (I) are 3D surface-rendered magnifications of the selected area above. (J, K) LC3 positive vesicles in unstimulated or treated BV2 (J) and primary microglial cells (K) were determined using ImageJ particle counting plugin after cell deconvolution (n=20). Results from at least three independent experiments were analysed by one-way ANOVA followed by Post-Hoc Dunnet’s test; n = 3. Error bars represent SEM (^∗∗∗^, P < 0.001), pMC: primary microglial cells.

We next aimed to determine the subcellular localization of fibrilar AS after cellular internalization and its distribution in comparison to the autophagy marker LC3. We stimulated BV2 cells stably expressing GFP-LC3 (BV2 GFP-LC3) with Alexa Fluor 594 AS fibrils for 12h, then, BV2 cells were stained for lysosomal-associated membrane protein 1 (LAMP-1). We found, by confocal microscopy and enhanced visualization by using 3D cell surface rendering approaches, that AS fibrils were confined to lysosomes and LC3 vesicles were distributed around them (FIG. 1D, F and G). In addition, we obtained similar results by using primary microglial cells (FIG. 1E, H and I). In order to confirm lysosomal localization for AS fibrils, we analysed the colocalization between the lysosomal specific enzyme N-acetylgalactosamine-6-sulfatase (GALNS) and fibrilar AS in primary microglial cells. Accordingly, these results showed high degree of colocalization between both labels (Fig. S2A).

In parallel experiments we studied the autophagic response by immunoblotting. Initiation of autophagy causes the conversion of LC3-I to LC3-II via the addition of a phosphatidylethanolamine (PE) group to the C terminus. We evaluated the conversion of LC3-I (non lipidated form with lower electrophoretic mobility) to LC3B-II (LC3 form C-terminally lipidated by PE, displaying higher electrophoretic mobility). In agreement with our previous results, we found an increase in the intensity of the LC3B-II band relative to the intensity of β-actin band after fibrilar but not monomeric AS stimulation (Fig. 1B and C). When Bafilomycin A1 (BAF) was added to fibrilar AS (hereafter fAS) stimulated cells, we observed an increase in the relative levels of LC3B-II in comparison with AS stimulation alone, indicating that fAS induces autophagy in microglial cells and it does not simply block autophagosome degradation (Fig. 1B and C). Overall, these results show that fAS has predominantly lysosomal localization after cellular internalization and it induces autophagy in microglial cells. The monomeric conformation was not able to activate the autophagy pathway at the time and dose studied.

### Autophagy dynamics of AS-stimulated microglial cells

To further study the autophagic response triggered by fAS in microglial cells, we conducted live-imaging experiments at different time points after fAS stimulation. We used BV2 GFP-LC3 microglial cells and LysoTracker Blue for lysosomal staining. We did not observe a significant increase in LC3 puncta during the first 8h after fAS stimulation (Fig. S1, SV6-9). Of note, fAS was quickly internalized during the first 20 min by microglial cells and it showed lysosomal localization since the earliest time points (Fig. S1C, SV5). Interestingly, after 12h of stimulation we detected a substantial increase in the autophagy response. LC3 vesicles increased over time and were predominantly associated with LysoTracker +/ fAS + vesicles forming a ring-like structure around them, as observed previously by confocal analysis (Fig. 2A and B, SV1 and SV2). In additional experiments, BV2 cells stably expressing GFP-ATG13 were co-transfected with CFP-LC3 plasmid. ATG13 integrates the autophagy initiation complex ULK1, the most upstream complex of the autophagy pathway and it is essential for autophagosome formation [5, 32]. In agreement with previous reports and autophagy dynamic studies, we observed positive ATG13 signal as an early event during autophagosome formation and its lifetime was shorter than the same structures containing LC3 (Fig. 2C and D, SV3 and SV4). As seen before, the LC3 signal associated with ATG13 progressed into characteristic rounded structures around synuclein fibrils.

**Figure 2.**
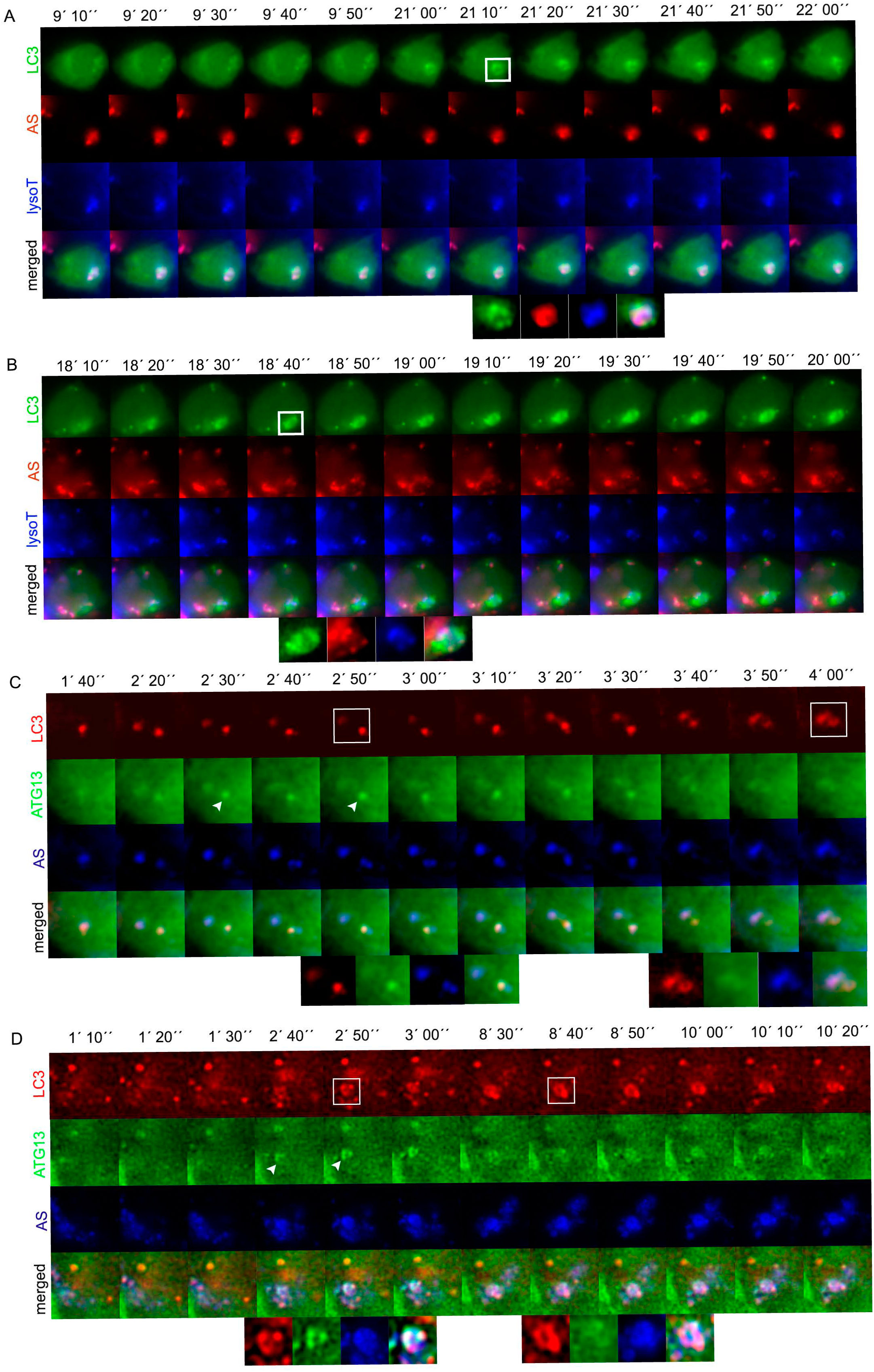
Evaluation of LC3 and ATG13 dynamics in AS-stimulated microglial cells by liveimaging. A, B. BV2 cells stably expressing GFP-LC3 (green) were stimulated with fAS (red) for 12h. Imaging was performed at 1 frame per 10 s during 1h and a selected interval within this sequence is shown. LysoTracker Blue was added 30 min previous to imaging acquisition. Note that autophagosomes form a ring-like structure around LysoTracker+/fAS+ structures. C, D. BV2 cells stably expressing GFP-ATG13 (green) were transfected with CFP-LC3 (red) and stimulated with fAS (blue) and imaged as described above. Arrowhead indicates the first discernible ATG13 punctum during autophagosome formation (C) and ATG13 forming a puncta pattern similar to LC3 (D).

Taken together these results indicate that the autophagic response to fAS follows a canonical route (utilising ATG13-positive structures that mature into LC3-positive structures), but it is not an immediate event after synuclein treatment. The facts that lysosomes containing fAS are surrounded by LC3 vesicles suggests the possibility that the autophagic machinery may respond to lysosomal damage caused by the fibrils, and this is a question we will address later.

### CLEM study of fAS-stimulated microglial cells evidences canonical autophagy

There are increasing reports showing the involvement of the non-canonical autophagy pathway in diverse pathological conditions since it was described for the first time. Although important advances have been made in the molecular characterization and differentiation between these alternative routes, we are still far from precisely understanding the mechanistic details and limits of both pathways [19, 46]. The principal difference between autophagosomes and non-canonical vacuoles is that the former have two limiting membranes positive for LC3 whereas the latter have one. In order to discriminate these different processes, we conducted CLEM experiments and analysed the presence of single or double membrane LC3-positive vesicles after fAS stimulation of BV2 GFP-LC3 microglial cells.

We clearly detected double membrane autophagic vesicles (AV) mainly correlating with LC3-GFP signal and closely associated with fAS+ structures (Fig. 3A-F, SV10, SV11 and SV12). Furthermore, we also observed double membrane vesicles and multi-membrane structures surrounded by a single-limiting membrane, probably as a result of fusion events between autophagosomes and lysosomes (Fig.3A-F, SV10, SV11 and SV12). These results are in agreement with a previous report describing similar AVs found on neocortical biopsies from AD human brain [51]. In agreement with Nixon et. al, the morphologies and composition of vesicles that accumulated after fAS treatment corresponded to those of the vesicular compartments of the autophagic pathway. Although we did not find enough evidence of quantitative ultrastructural analyses of glial organelles in the literature [7], most of the vesicles we observed in fAS-stimulated microglial cells correlated with standard morphometric criteria for the immature and mature autophagosomes as expected for neural cells [17, 18]. These criteria include a size >0.5 um in diameter, a double-limiting membrane (immature), and the presence within a single vesicle of multiple membranous domains from organelle sources such as, Golgi, mitochondria or endoplasmic reticulum (Fig. 3C, D and E, SV10, SV11 and SV12). We also observed similar AV morphology in non-treated microglial cells although single-membrane vesicles presented a smaller size in comparison to stimulated cells (Fig. S3). Moreover, in order to improve the detection of autophagosomes around fAS+ vesicles, we carried out additional CLEM experiments with fAS-stimulated microglial cells previously treated with BAF. Similar to our previous results, we found examples where an autophagosome was in close proximity to a larger AV (Fig. 3I and J) and as expected, we also detected areas where double-membrane autophagosomes accumulated around other AVs (Fig. 3 G and H, SV13).

**Figure 3.**
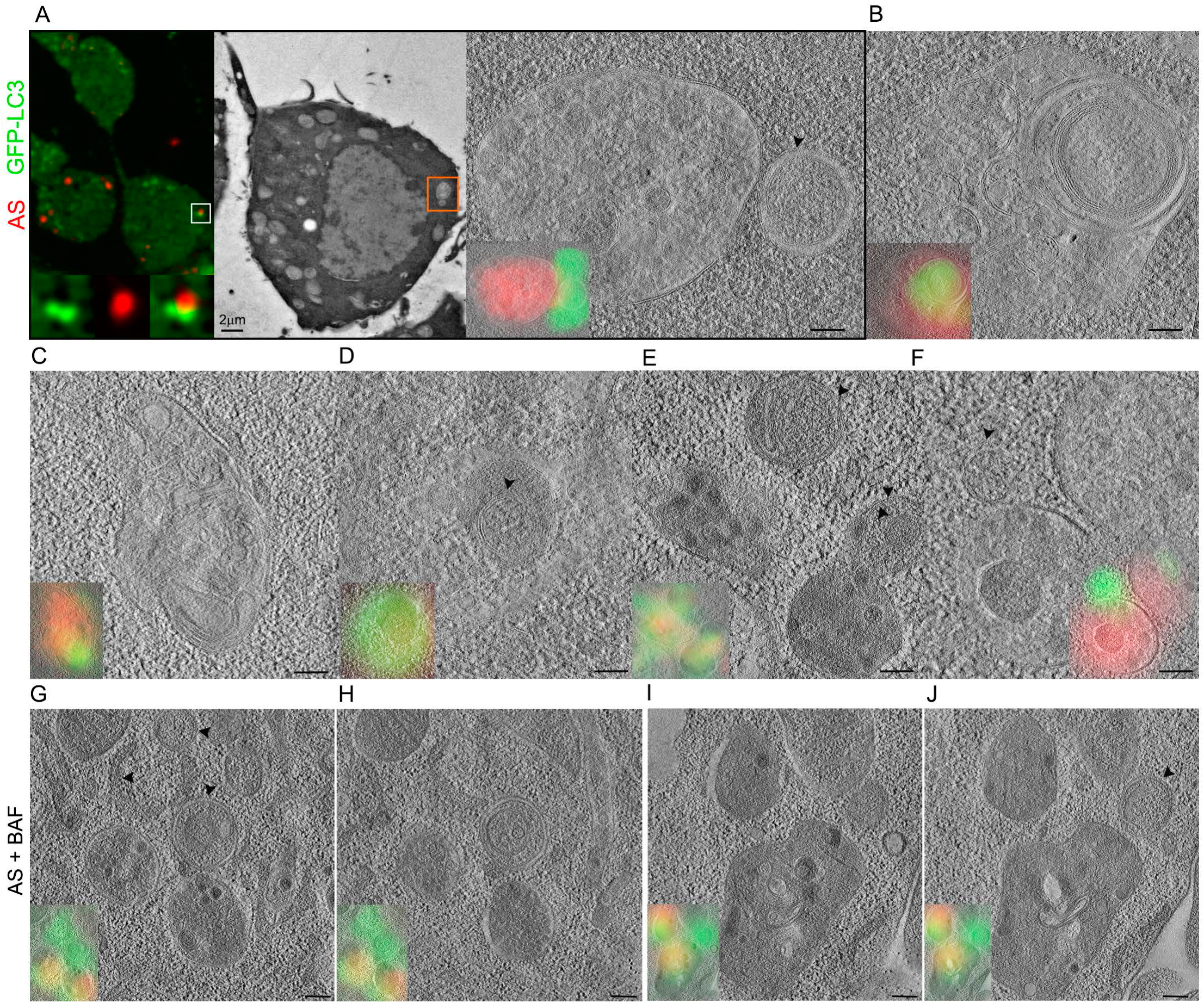
Correlative Light-Electron Microscopy study of LC3 positive vesicles in fAS-stimulated microglial cells. BV2 GFP-LC3 (green) cells were stimulated with fAS (red) for 12 h and incubated in the presence (G-J) or the absence (A-F) of BAF for the last 3h. Cells were then fixed by High Pressure Freezing (HPF) and processed for Optical and Electron Microscopy (EM) acquisition. 300 nm sections were imaged by fluorescence microscopy and EM tomograms were acquired at the regions of interest. (A) shows widefield and low (200X) and high-magnification (20000X) electron microscopy images with the overlay result. (B-J) show high-magnification slides (20000X) from representative tomograms of different cells indicating the overlay result (scale bar, 200nm). Arrowheads indicate double membrane vesicles.

To more precisely examine the ultrastructural nature of the AV formed after AS stimulation in microglial cells, we conducted live-CLEM assays. We stimulated BV2 GFP-LC3 cells with fAS for 12h and we followed the autophagy response by time-lapse widefield imaging and subsequent EM analysis (Fig. 4A and B). As we observed previously, an LC3 ring-like structure was formed around fAS (Fig. 4A). Interestingly, we found that the LC3-positive area predominantly correlated with a central double membrane autophagosome closely located to other single and double-membrane vesicles that were also positive for fAS signal (Fig. 4D and E, SV14 and SV15). In an additional live-CLEM experiment (Fig. 4C, G and H, SV16 and SV17) we observed that the region positive for LC3 signal correlated with concentrical multi-membrane structures with a central double membrane vesicle exhibiting a dense core. Moreover, these structures were in close contiguity with ER membranes, suggesting that the ER could be a main membrane donor in this process. These observations are in concordance with a previous electron tomography study showing that the ER associates with early autophagic structures and acts as a membrane source for autophagosome formation [28].

**Figure 4.**
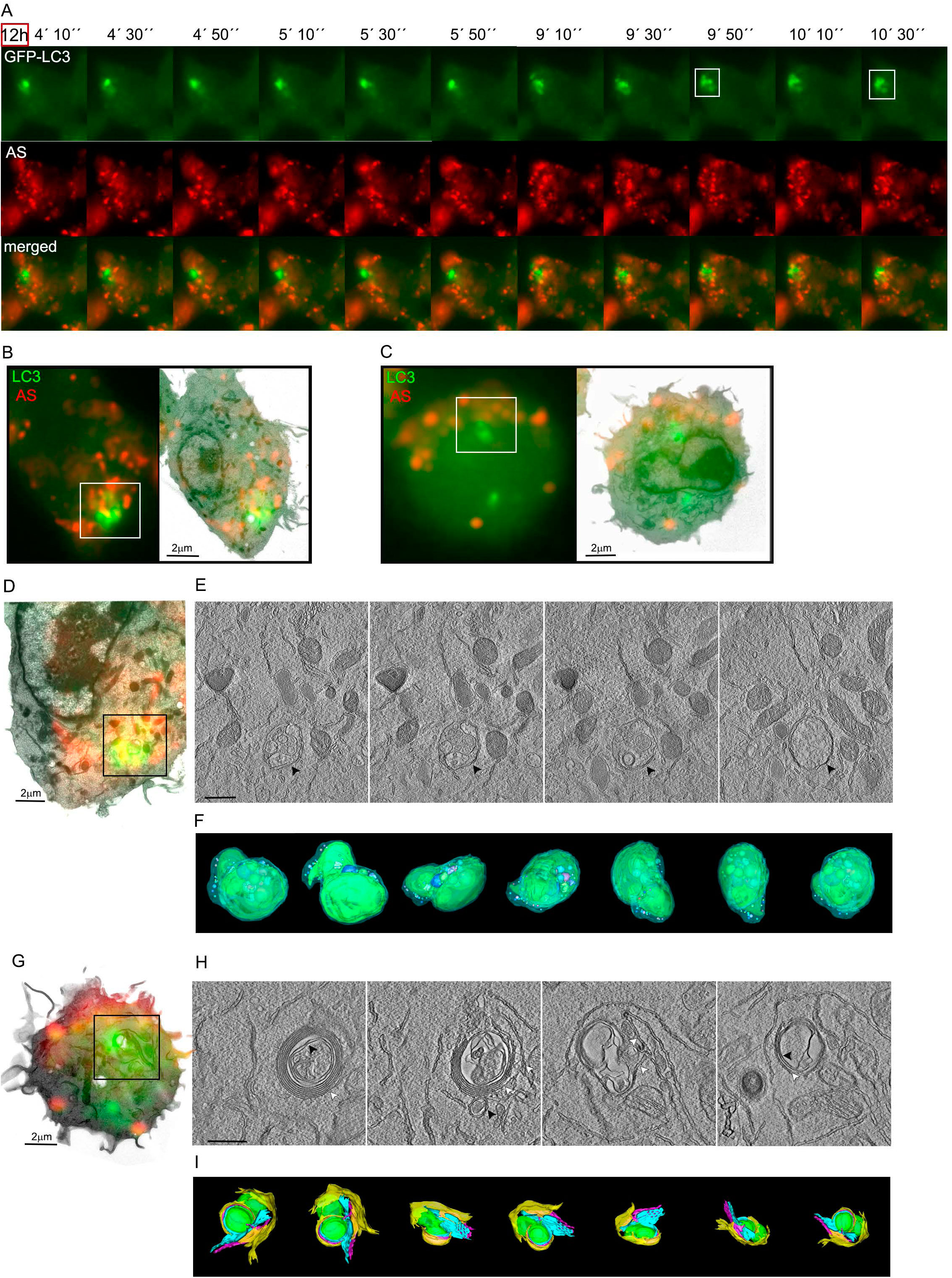
Live-CLEM imaging of fAS-stimulated microglial cells. (A) BV2 GFP-LC3 cells (green) were stimulated with fAS (red) for 12h. Imaging was performed at 1 frame per 20 s. After detecting the event of interest (white box), cells were immediately fixed and processed for EM tomography. Twenty serial tomograms were acquired on 300nm sections. (B) and (D) show the overlay result indicating the area acquired at high magnification (9400×) in each tomogram. (E) representative slides of the serial tomogram are shown (scale bar, 500nm). Arrowhead indicates a central double membrane autophagosome. (C, G). Correlation result of a second live-CLEM experiment indicating the area acquired at high-magnification (9400×) in each serial tomogram. BV2 GFP-LC3 cells were stimulated and imaged as described above. Representative tomogram slides are shown in (H). Black arrowheads indicate a double-membrane autophagosome surrounded by ER membranes (white arrows). (F) and (I) are 3D reconstructions of the vesicles shown in each serial tomogram’s slides.

Overall, these results provide evidence of canonical autophagy, instead of LAP, as the main effector pathway in microglial cells after fAS internalization.

### Effects of fAS on lysosomal and mitochondrial quality of microglial cells

Galectin-3 (Gal-3) is a sugar binding protein which recognizes beta-galactoside normally only present on the exterior leaflet of the plasma membrane and the interior leaflet of intracellular vesicles [60]. Gal-3 relocalization has been utilized to identify ruptured vesicles when bacteria and viruses enter the cytoplasm during infection [43, 62]. Recent studies have also demonstrated that even in the absence of bacteria or viruses, some galectins can translocate to damaged lysosomes before their removal by the autophagic pathway [24]. Since one of the possible mechanisms inducing autophagy in our cells may be following lysosomal damage, we used a recently described protocol to assess lysosomal damage recognised by Gal-3 [2]. We stimulated BV2 and primary microglial cells with labelled fAS at different time points and we analysed by confocal microscopy whether Gal-3 translocated to damaged lysosomes (Fig. 5 and Fig. S2D). L-leucyl-L-leucine methyl ester (LLOMe), which induces lysosome-specific membrane damage [63], was used as a positive control. In accordance with the autophagy dynamics previously described, we detected a significant change, from diffuse to punctate staining pattern, after 12h of fAS treatment but not during earlier time points (Fig. 5 and Fig. S2C). In addition, we observed a high extent of colocalization of Gal-3 with both fAS and Lamp-1 (Fig. 5C and D).

**Figure 5.**
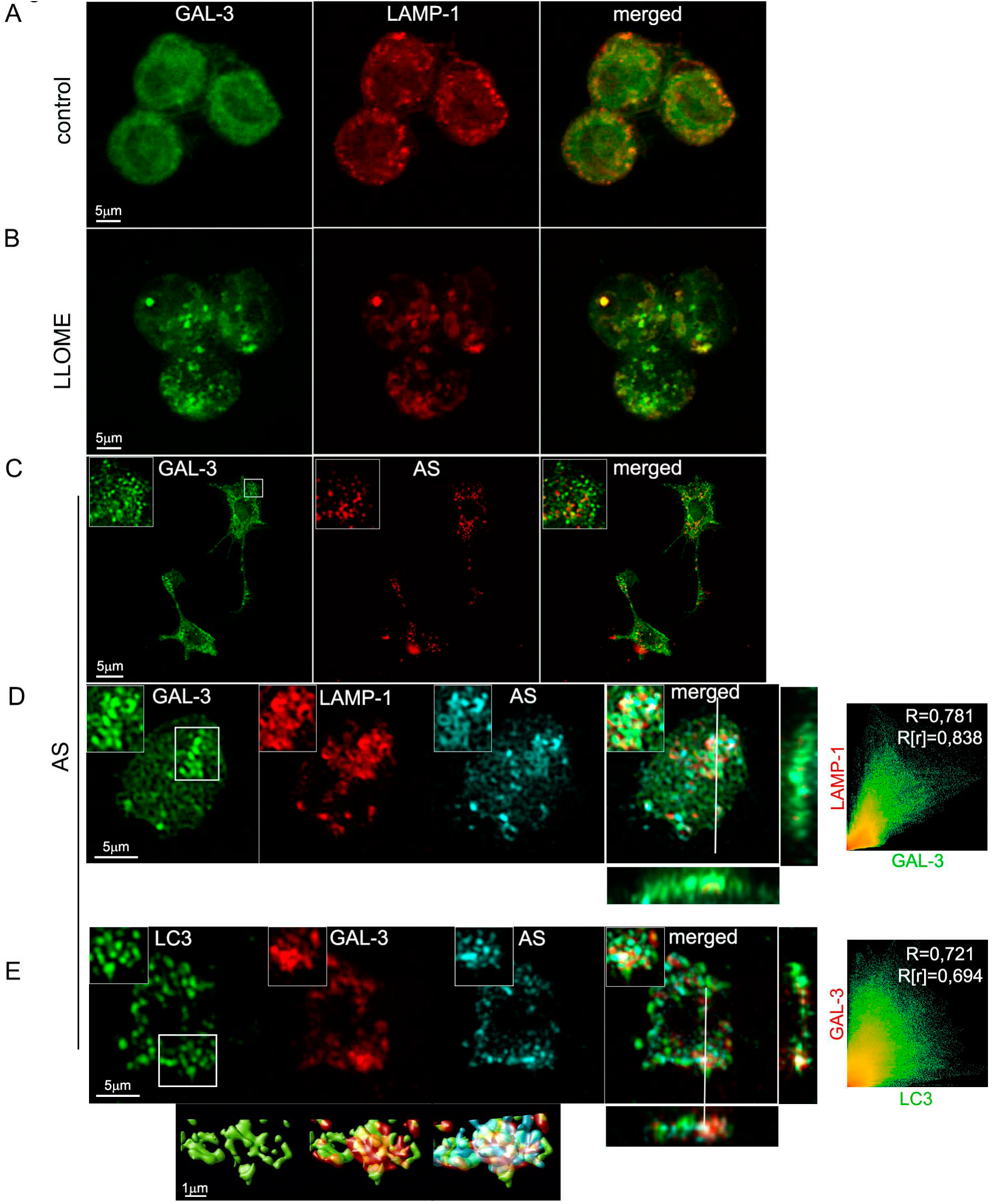
Evaluation of lysosomal damage in fAS-stimulated microglial cells. (A, B). BV2 cells were left untreated (A) or stimulated (B) with LLOME (500 μM) for 2h. After that, cells were immunostained with anti-Gal3 (green) and anti-Lamp1 (red) antibodies. (C, D, E) BV2 (C) or primary microglial cells (D, E) were treated with fAS for 12h. Cells were then fixed and stained for Gal-3 and Lamp-1 or LC3, as indicated. Merged images show orthogonal views and colocalization analysis between the specified labels. Pearson coefficient (R) and Overlap coefficient (R[r]) are listed.

We next evaluated if the LC3 positive vesicles formed after fAS stimulation colocalized with Gal-3 puncta and found a high degree of colocalization. This suggests that lysosomal damage acts as a positive signal for autophagy activation in fAS-stimulated microglial cells (Fig. 5E). These results are in agreement with the manuscript by Flavin et al. [21] describing that lysosomes ruptured by AS in SH-SY5Y neuronal cells are targeted for autophagic degradation. We also assessed if fAS could disturb lysosomal activity by using a Cathepsin-B fluorometric assay to monitor enzyme activity at different time points after microglial cell stimulation. We found a significant increase in Cathepsin-B activity after 8h of treatment which diminished to basal levels after 12h (Fig 6A), coincident with lysosomal impairment detection. These results indicate that lysosomes respond to the presence of fAS increasing Cathepsin-B activity which decreases with the progression of lysosomal damage.

**Figure 6.**
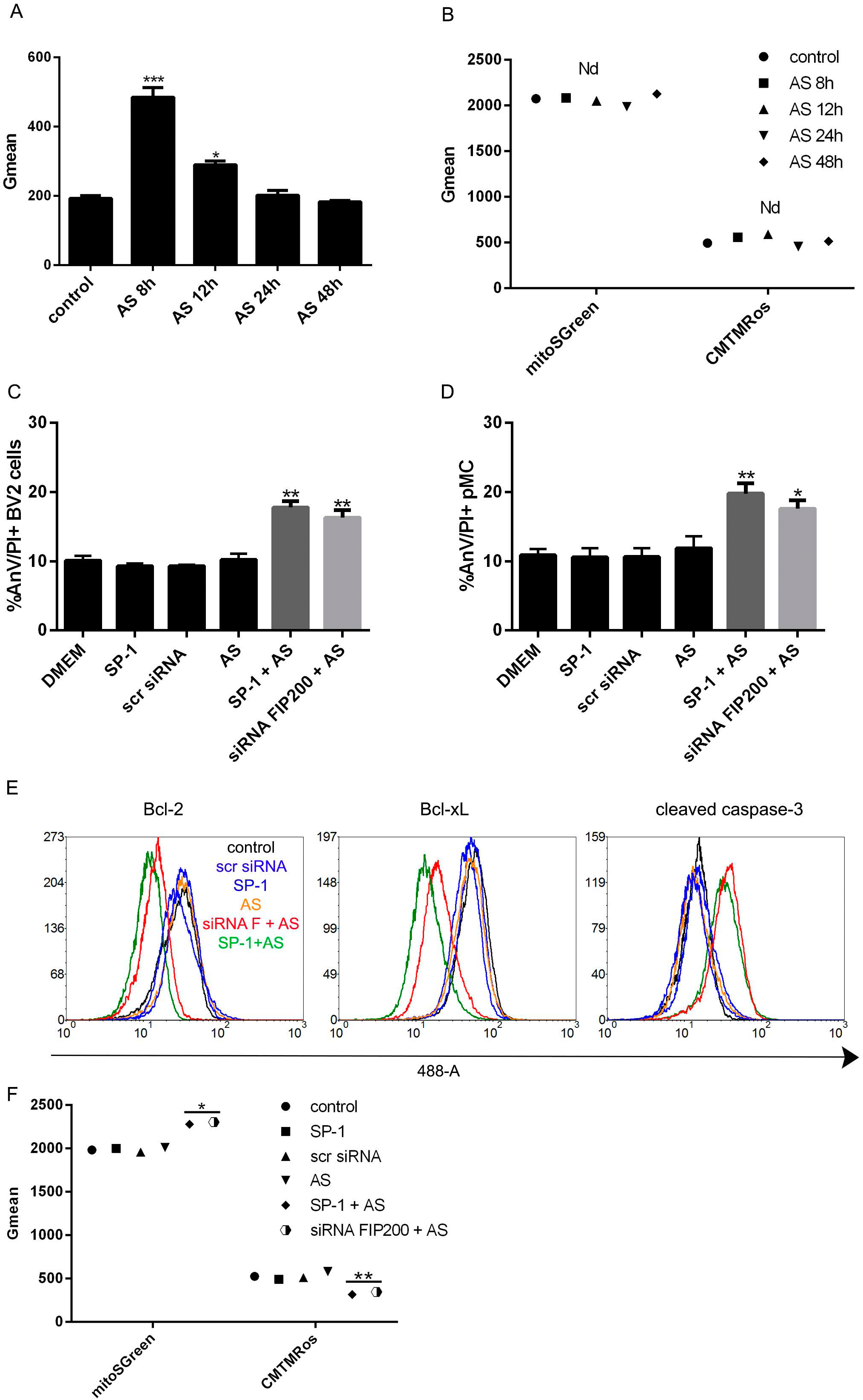
Effects of fAS stimulation on Cathepsin-B activity, mitochondrial quality and microglial cell survival. BV2 cells were stimulated with fAS at different time points. Microglial cells were then stained with MagicRed dye for evaluation of Cathepsin-B activity (A), MitoSpy Green FM for measuring mitochondrial mass or MitoSpy Orange CMTMRos for assessing mitochondrial membrane potential changes (B). Graphs show quantification of mean fluorescence intensity (Gmean) by flow cytometry. C, D. BV2 microglial cells (C) or primary microglial cells (D) were left untreated or treated with spautin-1 (10 μM) for 24h or FIP200 siRNA for 48h. Cells were then stimulated with fAS (5uM) for 24h and cell death was evaluated using propidium iodide (PI) combined with AnnexinV-FITC staining and subsequent flow cytometric analysis. Percentages of AnnexinV-FITC/IP double positive dead cells are shown. (E) BV2 microglial cells were treated and stimulated as described in (C) and Bcl-2, Bcl-xL and cleaved caspase-3 protein levels were evaluated by flow cytometry. Graphs show representative histograms for each protein. (F) BV2 microglial cells were left untreated or treated with spautin-1 (10 μM) for 24h or FIP200 siRNA for 48h and stimulated with fAS (5uM) for 24h. Mitochondrial mass and membrane potential were measured as described above (A, B). Results were analyzed by one-way ANOVA followed by Post-Hoc Dunnet’s test; n = 3. Error bars represent SEM (^∗^, P < 0.05; ^∗∗^, P < 0.01; ^∗∗∗^, P < 0.001).

In parallel experiments, we also evaluated mitochondrial status since mitochondrial dysfunction has been associated with several neurodegenerative diseases and AS was shown to alter mitochondrial activity [20]. However, we did not find changes either in mitochondrial mass nor mitochondrial cell membrane potential after fAS stimulation at the different time points studied (Fig. 6B). Overall these findings indicate that fAS induces lysosomal but not mitochondrial damage, with a similar kinetic as observed to autophagy activation, suggesting that this response is concomitant to lysosomal impairment.

### Autophagy prevents cell death in fAS-stimulated microglial cells

Autophagy is intimately associated with eukaryotic cell death and apoptosis. However the molecular connections between autophagy and cell death are complex and, in different contexts, autophagy may promote or inhibit cell death [4, 14, 25]. We therefore evaluated the effects of autophagy inhibition on microglial cell survival. We inhibited autophagy by using spautin-1, which promotes the degradation of VPS34 PI3 kinase complexes by inhibiting two ubiquitin-specific peptidases, USP10 and USP13 that target the Beclin1 subunit of VPS34 complexes [40]. In addition, we also used a siRNA targeting FIP200, a pivotal protein required for autophagy induction and autophagosome formation [27]. We evaluated the ability of spautin-1 and siRNA FIP200 to inhibit autophagy by confocal microscopy analysis of LC3 puncta formation after treating BV2 GFP-LC3 microglial cells with PP242, a specific mTORC1 inhibitor and autophagy inducer. We observed that both treatments significantly suppressed the autophagy response (Fig. S2B and C). Moreover, autophagy blockade by spautin-1 and siRNA FIP200 also decreased LC3 puncta formation in fAS-stimulated primary microglial cells (Fig. S2D).

We next investigated whether autophagy inhibition prior to fAS stimulation affects microglial cell viability. BV2 and primary microglial cells were cultured in the presence or the absence of spautin-1 or siRNA FIP200 and stimulated with fAS for 24h, then we evaluated microglial cell death by flow cytometry. We found an increase in the frequency of dead cells (AnV+/PI+) when autophagy was inhibited by these treatments in fAS-stimulated microglial cells. Similar results were obtained with both BV2 and primary microglial cells (Fig. 6C and D).

Mitochondrial outer membrane permeabilization (MOMP) is often required for activation of the caspase proteases that cause apoptotic cell death. As a consequence, mitochondrial outer membrane integrity is highly controlled, primarily through interactions between pro- and anti-apoptotic members of the B cell lymphoma 2 (BCL-2) protein family [61]. Bcl-2 and Bcl-xL anti-apoptotic proteins promote cell survival by preventing mitochondrial membrane permeabilization and subsequent content release which leads to caspase activation and ultimately, programmed cell death [59].

On the other hand, lysosomal damage and resulting lysosomal membrane permeabilization have been shown to induce apoptosis through MOMP, which can be brought about by cathepsin-mediated activating cleavage of pro-apoptotic Bid or inhibiting cleavage of anti-apoptotic Bcl-2 and Bcl-xL proteins [1, 12, 16]. Here, we also evaluated by flow cytometry the expression levels of Bcl-2, Bcl-xL and cleaved caspase-3 in fAS-stimulated BV2 microglial cells in the presence or the absence of both autophagy inhibitors. We found a decreased in Bcl-2 and Bcl-xL protein levels with a concomitant increase in cleaved-caspase-3 expression when autophagy was impaired (Fig. 6E). In parallel experiments we analysed mitochondrial mass and membrane potential changes after autophagy inhibition in fAS-stimulated microglial cells. We detected an increase in mitochondrial mass and a decrease in the mitochondrial membrane potential after autophagy blockade (Fig. 6F), which evidences the autophagy requirement for mitochondrial homeostasis. Collectively, our results showed that AS induced lysosomal damage and autophagy activation and the inhibition of this degradative pathway led to mitochondrial quality impairment, which includes MOMP, and consequent microglial cell death.

## DISCUSSION

Misfolding and intracellular aggregation of AS are thought to be crucial factors in the pathogenesis of Lewy body diseases (LBDs), such as PD. Recent studies suggest that small amounts of AS are released from neuronal cells by unconventional exocytosis, and that this extracellular AS contributes to the major pathological features of LBD, such as neurodegeneration, progressive spreading of AS pathology, and neuroinflammation [38]. In these neurodegenerative processes, the activation of microglia is a common pathological finding, which disturbs the homeostasis of the neuronal environment. Microglia’s behaviour is therefore a determinant on the disease’s progression.

In our present study, we show by confocal microscopy and immunoblotting analysis that fAS but not its monomeric conformation induces autophagy in microglial cells. Our results are in accordance with previous observations showing that AS fibrils are more potent cellular activators than other aggregation states. In agreement with a recent article from our group, Hoffman et al. found that fAS increased the production and secretion of pro-inflammatory cytokines in microglial cells in a greater extent than oligomers or monomers [9, 30].

Autophagy is a highly dynamic pathway and live-cell imaging has been extensively used to follow autophagy events in real-time [32]. Nonetheless, most of the studies monitoring the autophagic flux in glial cells were done by fluorescence microscopy of fixed cells and little is known about autophagy dynamics in microglial cells.

Here, we extensively characterized the autophagy dynamics of microglial cells stimulated with fAS by live-cell imaging. We observed that although fAS is quickly internalized by microglial cells, autophagy induction was evident after 12h of stimulation. Interestingly, we observed that LC3 decorated lysosomes containing fAS forming a ring-like structure around them. In additional experiments with stably expressing ATG13 BV2 cells we detected ATG13-positive structures that progresses into LC3-positive vesicles, which coincides with previous autophagy dynamic reports [32, 35] and suggest that the autophagic response to fAS follows a canonical pathway.

The kinetics of autophagy we observed contrasts with dynamics studies of LAP where phagosomes are rapidly decorated with LC3, usually within minutes [45]. Although we cannot rule out the involvement of the non-canonical pathway during this process, we could effectively correlate by high-precision and live-CLEM approaches, LC3-positive structures with double-membrane bound AV formed after fAS stimulation, indicating a predominant role for the canonical autophagy pathway rather than its alternative route during this process.

Recent evidence indicates that AS could disturb neuronal metabolism, by inducing lysosomal and mitochondrial damage [8, 24, 65]. Nevertheless, the effects of AS on glial organelles is poorly understood. Here, we report that fAS is incorporated into lysosomes after cellular internalization and it induces lysosomal damage in microglial cells at the same time point as when the autophagy response is significantly activated. Moreover, we showed a high degree of co-localization between Gal-3 puncta and LC3, suggesting that lysosomal damage rather than fAS acts as an activator signal for autophagy induction. In agreement with our results, Flavin et al. [21] showed that AS induced lysosomal rupture in SH-SY5Y neuroblastoma cells and LC3 colocalized with Gal-3-positive damaged vesicles. Moreover, the authors observed by immunofluorescence microscopic analysis on sections from PD patients that LBs aggregates were surrounded by Gal-3. Collectively, considering these literature evidences and our data shown here, we propose that accumulation of pathological protein aggregates and the formation of inclusions such as LBs arise from the failure of cellular attempts to degrade ruptured vesicles (and their amyloid contents) through the autophagy-lysosome pathway. Furthermore, Freeman et al. [24] showed that AS induces lysosomal rupture and Cathepsin-B release in neuronal cells after endocytosis. In addition, cysteine cathepsins were shown to be pivotal for lysosomal degradation of AS fibrils [49]. Although we observed a sharp increase in Cathepsin-B levels 8h after fAS stimulation, it notably diminished 12h after treatment which could indicate an overwhelmed lysosomal ability to degrade AS aggregates, leading ultimately to lysosomal damage. Although previous reports have indicated that AS induces mitochondrial damage in neurons [6, 44], we did not find significant changes either in mitochondrial mass or membrane potential at the different time points studied, indicating that fAS does not alter mitochondrial quality during the first 48h of stimulation in microglial cells.

There are increasing research articles providing evidence about the pivotal role of autophagy during CNS homeostasis and disease progression. The generation and analysis of the first nervous system-specific conditional knockout for ATG7, an E1-like enzyme that is essential for autophagy, revealed that while the conditional knockouts were born viable and were indistinguishable from control littermates for the first days of their life, they developed growth retardation as early as P14 and demonstrated various motor and behavioural deficits [33, 50]. Mice deficient for Atg5 specifically in neural cells have also been developed and analysed. These conditional mutants developed progressive deficits in motor function that are accompanied by the accumulation of cytoplasmic inclusion bodies in neurons. In Atg5−/− cells, diffuse, abnormal intracellular proteins accumulate and then form aggregates and inclusions [26]. Taken together, these evidences suggest that the continuous clearance of diffuse cytosolic proteins through basal autophagy is important for preventing the accumulation of abnormal proteins, which can disrupt neural function and ultimately lead to neurodegeneration.

Nonetheless, in the models discussed above autophagy was impaired not only in neurons but also in glial cells, yet the role of autophagy in glia and the contribution of defective glial autophagy in neurodegeneration remain poorly characterized.

In this report, we analysed the effect of disrupting autophagy on microglial cell survival after fAS stimulation. We observed by both spautin-1 treatment and down-regulation of FIP200 by siRNA increased levels of cell death of fAS-stimulated microglial cells. Our findings agreed with a previous report indicating that neural-specific deletion of FIP200, involved in autophagosome biogenesis, caused axonal degeneration in cerebellar neurons eventually causing their death [39].

The mitochondria-mediated caspase activation pathway is a major apoptotic pathway characterized by MOMP and subsequent release of cytochrome c into the cytoplasm to activate caspases. MOMP is regulated by the Bcl-2 family of proteins which act as inducers or blockers of the process [66]. Of relevance to this work, it has been shown that lysosomal damage can induce MOMP-dependent cell death and lysosomal proteases released into the cytosol have been implicated in apoptotic cell death [57]. In the present study, we found that autophagy inhibition increased mitochondrial mass but impaired mitochondrial membrane potential, down-regulated Bcl-2 and Bcl-xL protein levels and increased cleaved caspase-3 protein expression in fAS-stimulated BV2 cells, which suggest activation of apoptosis. Although we cannot disregard additional upstream signals triggering MOMP, we consider it likely that lysosomal damage, together with the inability of the autophagic pathway to clear this organelle, leads to MOMP and ultimately to cell death. Collectively, our results suggest a protective role for the autophagy pathway in fAS-stimulated microglial cells.

Extracellular AS, has emerged as a crucial player in the pathogenesis of LBDs and, possibly, MSA. Recent studies have provided evidence that extracellular AS alone, particularly fibrils, can be responsible for all the major pathological changes in neurodegenerative diseases: aggregate deposition and spreading, neuroinflammation and neurodegeneration [55]. Several articles have indicated that extracellular AS activates microglial cells producing an increase in the release and production of pro-inflammatory mediators. However, this is to the best of our knowledge, the first study showing that fAS induces lysosomal damage and autophagy in microglial cells, describing the dynamics of this response and correlating light-microscopy imaging with the specific subcellular autophagic compartments participating during this process. In this manuscript we provide new insights into the effects of AS on microglial autophagy and cell survival and we propose that AS-induced lysosomal damage activates canonical autophagy as a rescue mechanism. Future research evaluating the molecular mechanisms triggered by protein aggregates, such as AS, on glial cells would shed light on novel therapeutic targets for neurodegenerative disorders.

## Acknowledgements

This work was supported in part by FONCyT, CONICET and SECyT-UNC, Argentina. Its contents are solely the responsibility of the authors and do not necessarily represent the official views of CONICET. C.B. thanks to EMBO, EMBL, Boehringer Ingelheim Fonds, IUBMB Wood-Whelan Fellowship and to The Company of Biologists Travelling Fellowships for supporting short-research stays abroad.

## Author Contributions

C.B. performed all experiments, designed the research study and wrote the manuscript. J.M.P.R. and D.S.A. collaborated in cell culture, westernblot, confocal microscopy and flow cytometry experiments. P.R and A.K collaborated in the design, acquisition and analysis of CLEM experiments, O.F. contributed with BV2 GFP-LC3 production and edited the manuscript, J.I.G. and M.S.C. contributed reagents/materials/analysis tools, Y.S. contributed with the design of CLEM experiments and edited the manuscript, N.T.K. contributed with reagents/materials/analysis tools, the design of the study and edited the manuscript, P.I. designed the research study and wrote the manuscript and is the corresponding author and holds all the responsibilities related to this manuscript.

---

**Figure S1.**
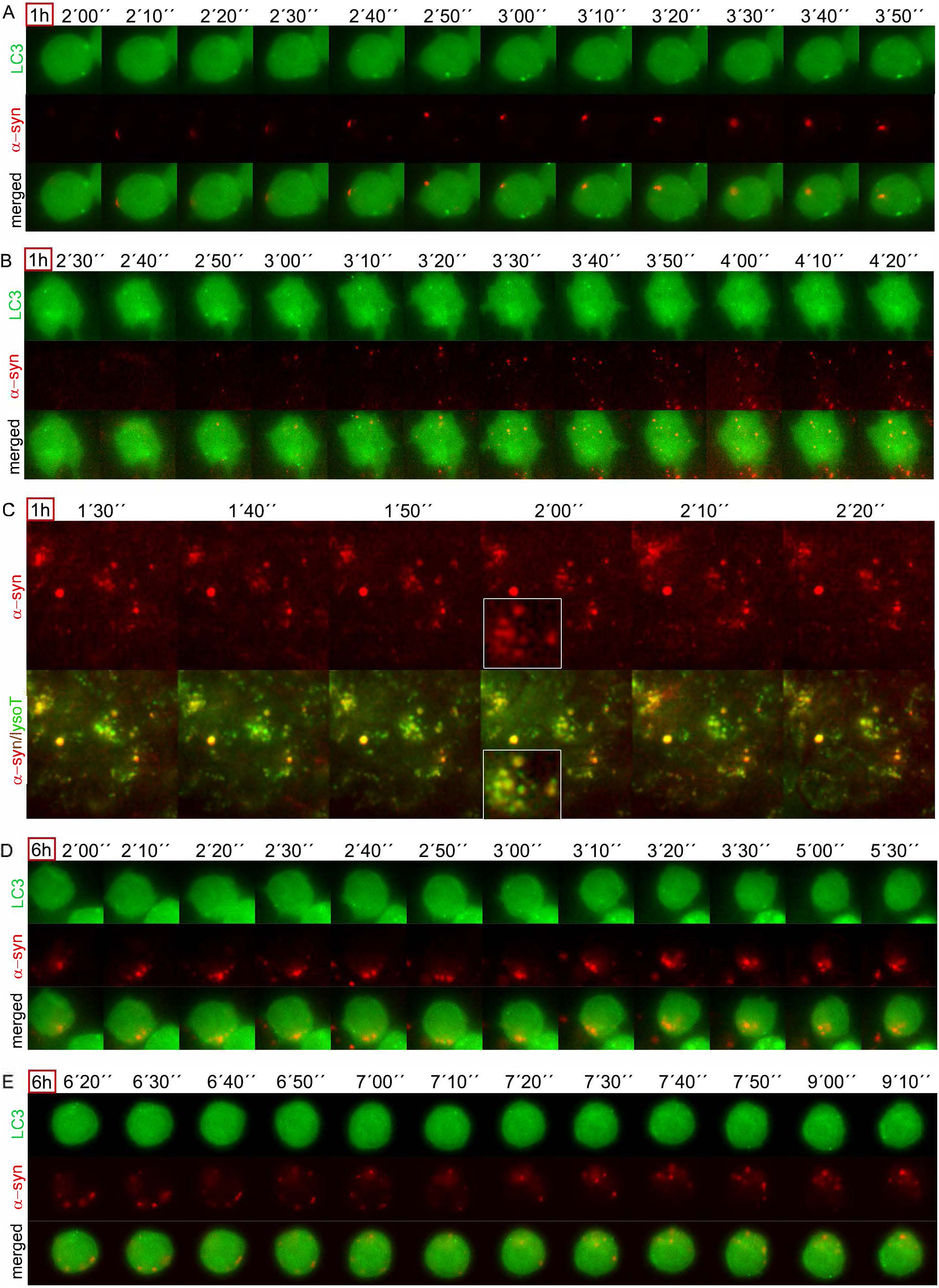
Autophagy dynamics of fAS-stimulated BV2 GFP-LC3 cells during early time points. A, B, C. BV2 GFP-LC3 (green) cells were stimulated with fAS (5uM, red) for 1h. Imaging started immediately after cellular stimulation and it was performed at 1 frame per 10 s during 1h, a selected interval within this sequence is shown. (C) shows fAS (red) and LysoTracker staining (green) of BV2 GFP-LC3 cells stimulated and imaged as described above. Of note that synuclein/LysoTracker colocalization is quickly observable after cellular internalization. D, E. BV2 GFP-LC3 (green) cells were stimulated with fAS (red) for 6h and imaged at 1 frame per 10 s during 1h, a selected interval within this sequence is shown.

**Figure S2.**
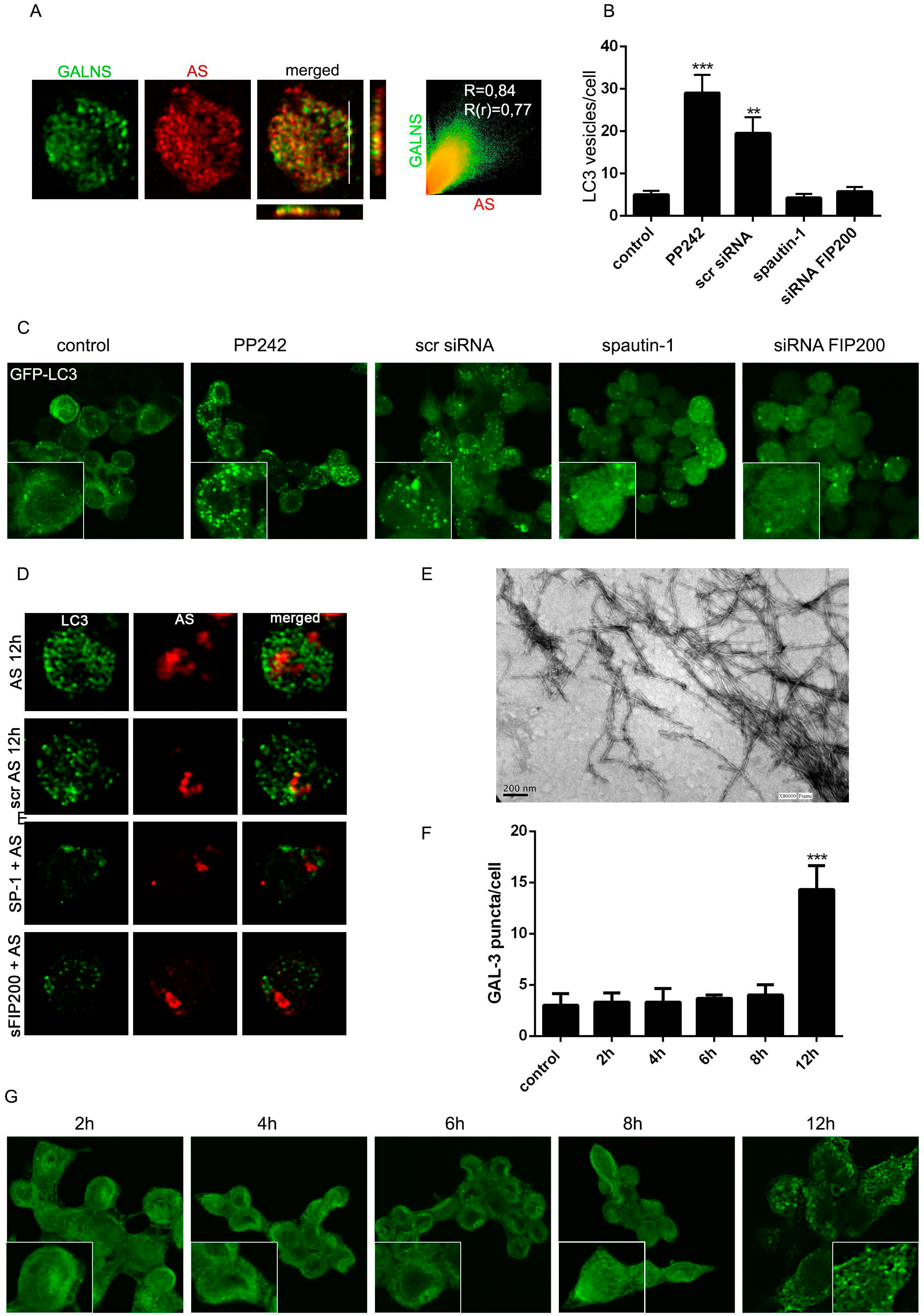
(A) Primary microglial cells were stimulated with fAS for 12h and co-localization between lysosomal specific marker GALNS and fAS was analysed by confocal microscopy. Pearson coefficient (R) and Overlap coefficient (R[r]) are listed. (B) LC3 positive vesicles in BV2 GFP-LC3 cells treated with the mTOR inhibitor PP242 (100uM) for 3h in the presence or the absence of spautin-1 (10 μM) or siRNA FIP200 were determined using ImageJ particle counting plugin. Spautin-1 and siRNA FIP200 were added 24h and 48 h before autophagy stimulation, respectively. Control treatments were used in the same conditions. (C) Representative Immunofluorescence images of BV2 GFP-LC3 treated as described in B. (D) primary microglial cells were cultured in the presence or the absence of spautin-1 (10 μM) or siRNA FIP200 and stimulated with fAS (red, 5uM). After 12h cells were immunostained for LC3 (green) and confocal images after cellular deconvolution are shown. (E) TEM of fAS (scale bar, 200nm). (F, G) BV2 microglial cells were stimulated with fAS at the indicated time points. After that, cells were immunostained for GAL-3 and puncta formation were determined using ImageJ particle counting plugin (E). Results were analysed by one-way ANOVA followed by Post-Hoc Dunnet’s test; n = 3. Error bars represent SEM (^∗∗^, P < 0.01; ^∗∗∗^, P < 0.001).

**Figure S3.**
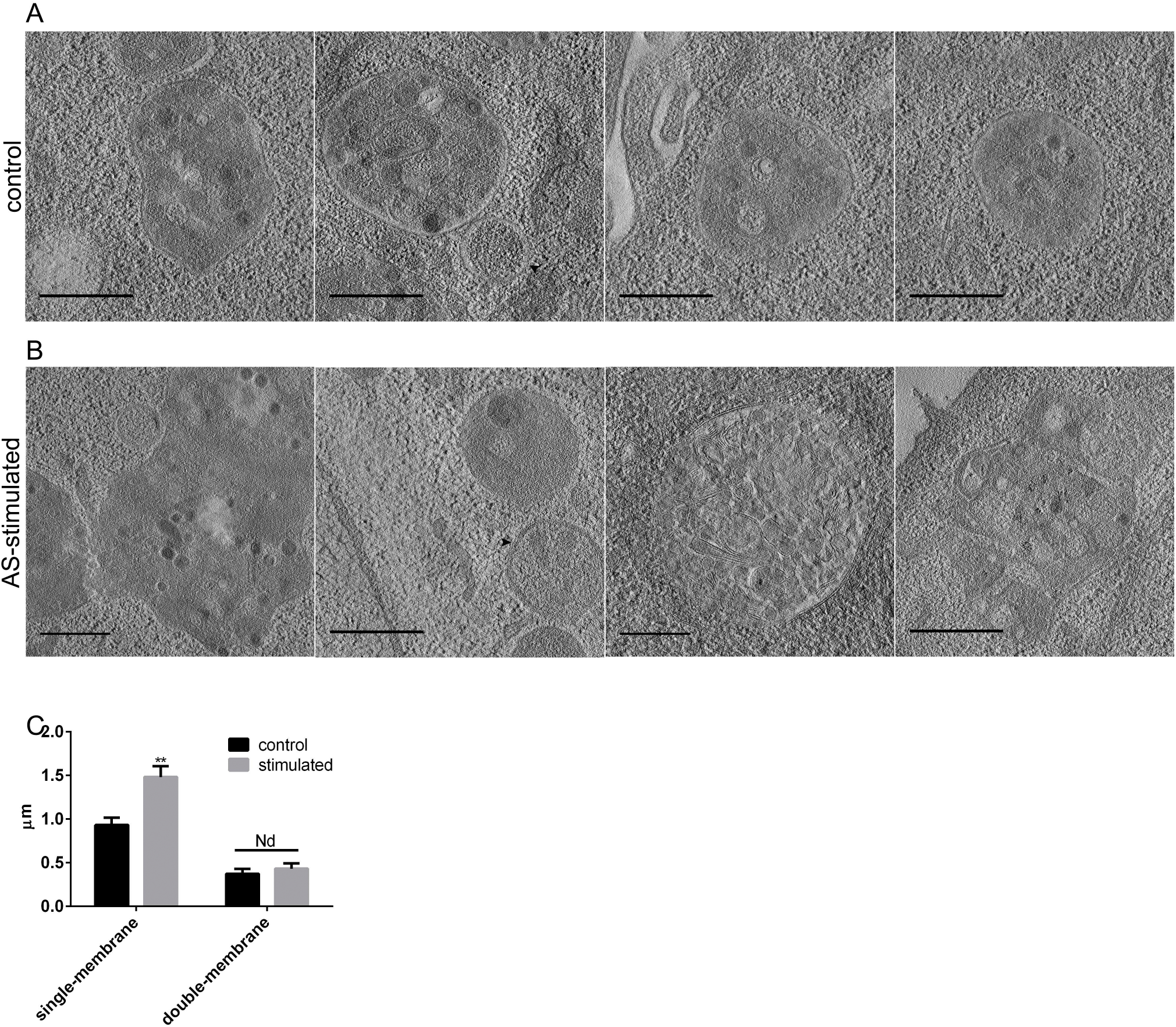
Comparison of single and double-membrane vesicles found in control and fAS-stimulated BV2 microglial cells. A, B. Representative images from EM tomograms of control (A) and fAS-stimulated (B) BV2 GFP-LC3 microglial cells showing different single and double membrane AV (scale bar, 500nm). (C) Graphs show the AV size quantification in both conditions. Results were analyzed by one-way ANOVA followed by Post-Hoc Dunnet’s test; n = 30. Error bars represent SEM (^∗^, P < 0.05; ^∗∗^, P < 0.01).

## Supplementary Videos

(SV1-SV9) Video files showing autophagy dynamics at different time points after fAS stimulation of BV2 microglial cells.

SV1, SV2. BV2 GFP-LC3 cells were stimulated with fAS for 12h and imaged at 1 frame per 10 s during 1h. SV1 and SV2 correspond to the sequence shown in Fig. 2A and Fig. 2B, respectively. Of note that LC3 (green) forms a ring-like structure around fAS (red). LysoTracker (blue) was used for lysosomal staining.

SV3, SV4. BV2 cells stably expressing ATG13 (green) were co-transfected with CFP-LC3 plasmid (red). Microglial cells were stimulated with fAS (blue) for 12h and imaged at 1 frame per 10 s during 1h. SV3 and SV4 correspond to sequence shown in Fig. 2C and Fig. 2D, respectively. Of note that ATG13-positive structures mature into LC3-positive vesicles. SV5. BV2 GFP-LC3 cells were stimulated with fAS (red) and imaged immediately after stimulation for 1h. LysoTracker (green) was used for lysosomal staining.

SV6, SV7, SV8, SV9. BV2 GFP-LC3 cells (green) were stimulated with fAS (red) and imaged immediately (SV6 and SV7) or 6h after stimulation (SV8, SV9). Of note that no significant change in the dynamics of autophagy is detected over time, unlike long-term stimulation.

SV10 – SV13. EM tomograms from fAS-stimulated microglial cells.

EM high-magnification (20000X) tomograms corresponding to the images indicated as Fig. 3A (SV10), Fig. 3B (SV11), Fig. 3E (SV12) and Fig. 3G (SV13) are shown as video files.

Of note, the presence of double membrane autophagosomes.

SV14, SV15. Live imaging video file (SV14) and serial tomogram and 3D model (SV15) corresponding to Fig 4A and Fig. 4E, respectively. Tomograms of twenty consecutive 300nm sections were acquired at 9400X magnification and later aligned. SV15 shows a central double-membrane autophagosome containing several vesicles.

SV16, SV17. Live imaging video file (SV16) and serial tomogram and 3D model (SV17) corresponding to Fig 4C and Fig. 4H, respectively. SV17 shows an inner autophagosome and multiple ER membranes surrounding it, suggesting it is an immature AV. Of note, an additional concentrical structure exhibiting a dense core is also exhibited.

